# Integrating promiscuous enzyme activities in protein-constrained models pinpoints the role of underground metabolism in robustness of metabolic phenotypes

**DOI:** 10.1101/2024.09.06.611666

**Authors:** Maurício Alexander de Moura Ferreira, Eduardo Luís Menezes de Almeida, Wendel Batista da Silveira, Zoran Nikoloski

**Affiliations:** Department of Microbiology, Federal University of Viçosa, Viçosa, Minas Gerais, 36570900, Brazil; Bioinformatics, Institute of Biochemistry and Biology, University of Potsdam, Potsdam, 14476, Germany; Systems Biology and Mathematical Modelling, Max Planck Institute of Molecular Plant Physiology, Potsdam, 14476, Germany

## Abstract

The integration of enzyme parameters in constraint-based models have significantly improved the prediction of physiological and molecular traits, including enzyme resource usage and distribution. However, current approaches largely neglect the set of promiscuous enzyme activities that jointly comprise the so-called underground metabolism. To allow enzyme-constrained study of underground metabolism, we developed the CORAL Toolbox. This toolbox reworks enzyme usage into subpools for each reaction catalysed by a promiscuous enzyme, increasing the resolution of modelled enzyme resource allocation. Applying CORAL with an enzyme-constrained genome-scale metabolic model of *Escherichia coli*, we found that underground metabolism resulted in larger flexibility in metabolic fluxes and enzyme usage. Knocking out the main activity of a promiscuous enzyme led to small enzyme redistribution to the side activities. Further, knocking out pairs of main activities showed that non-promiscuous enzymes exhibited larger effect on growth. In addition, we demonstrated these findings are robust with respect to the parameterization of the models with catalytic rates from different prediction tools. Together, our results from modelling underground metabolism in enzyme-constrained models indicated that promiscuous enzyme activities are vital to maintain robust metabolic function and growth.

## 1 Introduction

Enzymes are the workhorses of metabolism. They catalyse the conversion of substrates into products for the majority of metabolic reactions. Although some enzymes are highly specific with respect to reactions they catalyse, there is notable enzyme promiscuity, with estimates of some enzymes catalysing reactions with hundreds of different metabolites serving as substrates ^1^. Promiscuous enzymes catalyse more than one reaction by binding with smaller affinity to other substrates ^2^. As a consequence of lower substrate affinity, the side reactions catalysed by promiscuous enzymes occur at a lower rate than the main reaction. Promiscuous enzymes also exhibit lower catalytic efficiency for the side reaction in comparison to the main reaction ^3^. As a result, it has been taught that side reactions are physiologically irrelevant ^4^. However, these side reactions still happen often enough to form an alternative metabolic network, termed underground metabolism ^5^. This underground metabolic network serves an important role in evolution, providing the reservoir of enzyme functions to evolve via natural selection ^6^. Evidence indicates that gene duplication events contribute to underground metabolism by creating copies of the enzyme, with one copy evolving higher affinity to the substrate of a certain side reaction ^7^. Further, underground metabolism can also be used for biotechnological purposes, aiding laboratory adaptive evolution (ALE) experiments ^8^ and guiding metabolic engineering efforts ^9^.

Constraint-based approaches rely on optimization principles to predict and study metabolic phenotypes using genome-scale metabolic models (GEMs) ^10^. GEM-based investigations of underground metabolism have provided useful insights, such as the connectivity between native and underground metabolism and how underground reactions contributes to adaptation to new environments ^6^, improved gap-filling of GEMs ^11^, and the design of metabolic engineering strategies ^9^. While insightful, these investigations were performed using conventional GEMs, which do not consider constraints on the available enzyme abundance and enzyme catalytic rates. Consideration of these constraints has resulted in the generation of protein-constrained GEMs (pcGEMs) that have been shown to improve predictive performance ^12–14^. However, it remains elusive how the metabolic network allocates enzyme resources between main and side reactions.

The GECKO ^13,15,16^ formulation of pcGEMs directly represents enzymes in the stoichiometric matrix, where each enzyme corresponds to a pseudometabolite that participates in a reaction, with a pseudo-stoichiometric coefficient given by the ratio between the molecular weight of the enzyme (MW) and its turnover number, *k_cat_*, for the reaction. Therefore, the GECKO formulation of pcGEMs allows for the simultaneous prediction of the enzyme usage distribution vector alongside a flux distribution using conventional constraint-based approaches, like flux balance analysis (FBA). The enzyme usage distribution is bounded by the total protein pool. The total protein pool is represented as an exchange pseudoreaction, itself bounded by experimental data on total cell protein content, and for each enzyme there is a pseudoreaction assigning a share of the total protein pool to the enzyme. In this formulation, an enzyme that catalyses two or more reactions uses the same enzyme pseudometabolite in each reaction, indicating that the same amount of enzyme is used for all reactions catalysed by the enzyme. However, since most enzymes are likely occupied by their main substrates, the availability of enzyme resources to side reactions may be drastically reduced. This issue is also not accounted in the existing pcGEM approaches.

To address these questions, here we propose an approach for modelling promiscuous enzyme activity in pcGEMs, termed **CO**nstraint-based p**R**omiscuous enzyme **A**nd underground metabolism mode**L**ling (CORAL) Toolbox. The CORAL Toolbox builds on GECKO 3 to predict enzyme allocation while ensuring that a subpool allocation to a promiscuous enzyme is used for a particular reaction. We show that CORAL can predict the distribution of resources among main and side reactions, making it a useful approach to understand promiscuous enzyme activity and underground metabolism. In addition, we show how CORAL can be used to gain insights in the functional implications of promiscuous activities in metabolic networks.

## 2 Material and methods

### 2.1 Refining the iML1515 model and reconstructing the pcGEM

To reconstruct a GEM accounting for underground metabolism, we manually curated the *Escherichia coli* GEM iML1515 ^17^ to include underground reactions (see Supplementary Data 1). We included the reactions from the underground iJO1366 ^18^ model created in the Kóvacs et al. (2022) study ^9^. These were predicted by the PROPER algorithm, with 20% of these reactions validated experimentally ^6,19^. We integrated these reactions, matched all annotations from iJO1366 to the format used by iML1515, and standardized the gene-protein-reaction (GPR) rules, resulting in the iML1515u model.

Next, we reconstructed the protein-constrained version of iML1515u using GECKO 3 ^15^. We populated the adapter file with *E. coli*-specific information, using a maximum growth rate when not constrained by nutrient uptake of 0.6 h^-1^ ^20^, total protein content (*P_tot_*) of 0.61 g_protein_/g_DW_^21^, and the default value of 0.5 for average enzyme saturation factor (*σ*) and fraction of enzymes in the model (*f*). We followed the steps described to reconstruct the full ecModel, with constraints on individual enzymes. For all reactions in the model, we integrated *k_cat_* values predicted by DLKcat ^22^. We also built a model with *k_cat_* values predicted by TurNuP ^23^ for underground reactions to assess how a different *k_cat_* prediction tool would affect the results. We followed the protocol until Stage 2 since Stage 3 deal with tuning model parameters as we wanted to preserve the original DLKcat-predicted *k_cat_* values. All model refinements were performed using the COBRA Toolbox 3 ^24^ and the RAVEN Toolbox 2 ^25^ in MATLAB (The MathWorks Inc., Natick, Massachusetts).

### 2.3 Restructuring the pcGEM to account for enzyme usage in underground metabolism

To account for the enzyme usage for individual reactions catalysed by a promiscuous enzyme, we developed the CORAL Toolbox. CORAL modifies how enzymes are used by splitting the pool of an enzyme that catalyses more than one reaction into multiple subpools, with each subpool responsible for the enzyme resources of only one reaction (Figure 1). The restructuring of a GECKO 3-constructed pcGEM is done in three steps: (i) simplifying GPR rules; (ii) splitting enzyme pools into subpools for all reactions; and (iii) updating enzyme information on the model.

**Figure 1.**
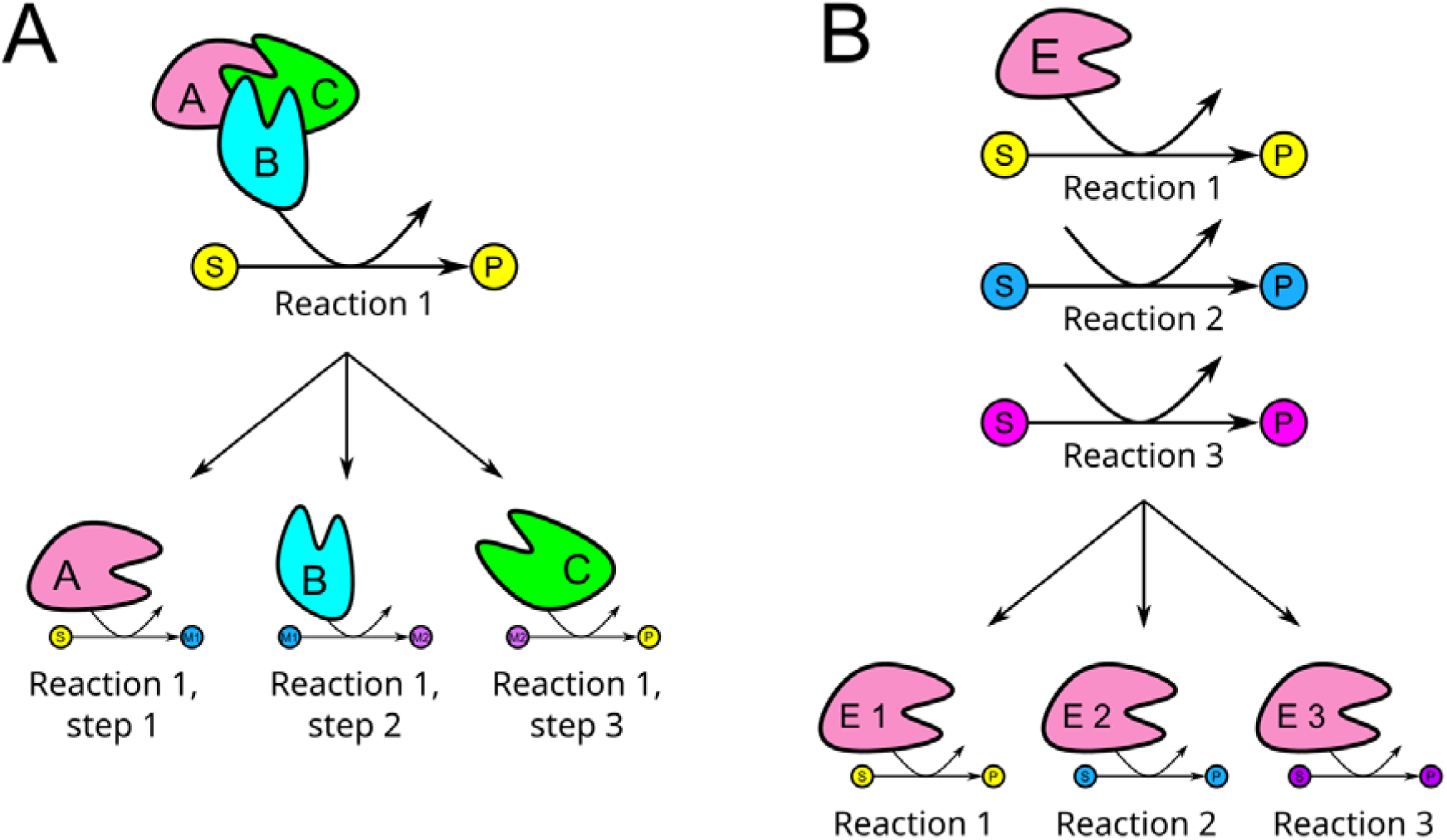
Restructuring of the pcGEM as performed in CORAL. **A)** Simplification of GPR rules to split reactions catalysed by enzyme complexes into multiple partial reactions. **B)** Splitting of promiscuous enzymes into subpools of enzymes, each catalysing a single reaction.

The first step of the restructuring entails the simplification of the GPR rules. In GECKO 3, reactions catalysed by isozymes (“OR” rules) are separated into different reactions catalysed by a single enzyme. The CORAL Toolbox introduces another simplification, dealing with reactions catalysed by multiple enzymes (“AND” rules), by splitting all reactions catalysed by enzyme complexes into multiple partial reactions catalysed by one enzyme each.

In the second step, we ordered the reactions catalysed by promiscuous enzymes from the lowest to the highest ratio of molecular weight (MW) to turnover number, i.e. MW/*k_cat_*. We defined as the main reaction the one with the largest *k_cat_* value (i.e., lowest MW/*k_cat_* ratio); the other reactions catalysed by the same enzyme are considered side reactions. For each reaction catalysed by a promiscuous enzyme, a pseudoenzyme was created to replace the promiscuous enzyme. A number was then appended to the IDs of each pseudoenzyme according to the order of MW/*k_cat_* ratios. These pseudoenzymes comprises a subpool that draw resources from the enzyme pool, specific to one enzyme. Each enzyme pool then draws resources from the total protein pool, which is obtained from an exchange reaction.

Lastly, GECKO 3 introduces a new structure in the YAML and MAT file formats, the “.ec” structure, where all enzyme information is stored. Given that CORAL changes how enzymes are integrated in the model, most of this information no longer matches what is present in the model after the changes take place. The third step updates all enzyme information and makes it available at the new “.und” structure, while the original “.ec” structure remains available.

### 2.4 Flux variability analysis

To evaluate the ranges of metabolic fluxes and of enzyme usages in the CORAL model, we performed flux variability analysis (FVA). We first maximized and minimized the flux *v* of reaction *j* to find its flux range:

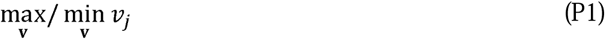

subject to:

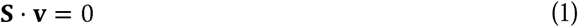

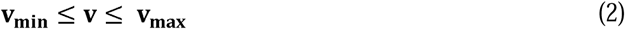

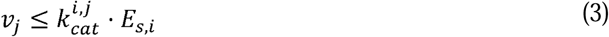

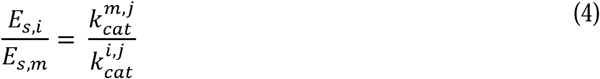

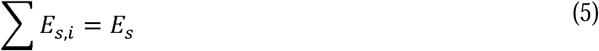

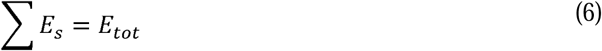

where **S** is the stoichiometric matrix, **v** is the flux distribution vector, **v_min_** is the flux lower bound vector, **v_max_** is the flux upper bound vector, 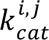 is the *k_cat_* value for the enzyme subpool *i*, catalysing the reaction *j*, *E* is the enzyme usage in the enzyme *s*, subpool *i*, *E_s,m_* is the enzyme usage for any subpool *m* that is not catalysing the main reaction, 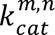 is the *k_cat_* value for enzyme *m*, and *E_tot_* is the total protein pool. The ratio constraint introduces a proportional relationship where more inefficient enzymes (i.e., lower *k_cat_* values) would require a higher allocation of enzyme resources to carry the same amount of flux. In other words, we aim to model the prioritization of enzyme resources based on catalytic efficiency of the enzyme. Next, we performed FVA on the enzyme usage pseudoreactions using the same constraints as before, but with the following objective function:

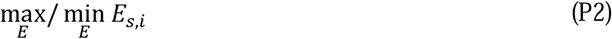

### 2.5 Single and double knockouts of subpools

We investigated how enzyme resources are distributed to side reactions when the main reaction is knocked out. To this end, we first used a modified implementation of parsimonious FBA (pFBA), whereby we minimize the sum of enzyme subpools instead of sum of fluxes. In the first round we obtain the enzyme subpool distribution we later use in a second round:

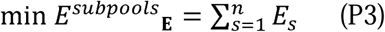

subject to:

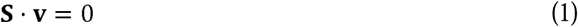

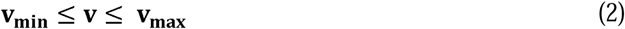

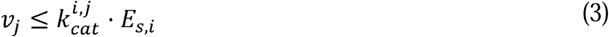

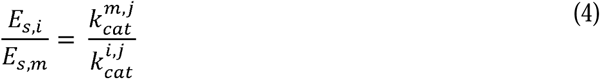

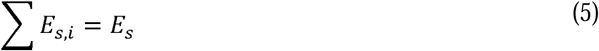

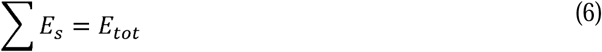

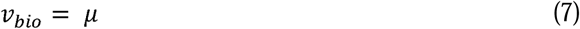

where *n* is the number of enzymes, *v_bio_* is the flux through the biomass pseudoreaction, and *μ* is the specific growth rate.

After finding the enzyme subpool distribution, it is next used to constrain the model (allowing 5% flexibility) in a second round of simulation, where we set to zero the subpool *E*_*s*,1_ of each enzyme *s* one by one, and optimize growth:

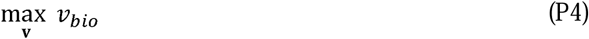

subject to:

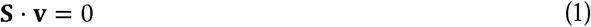

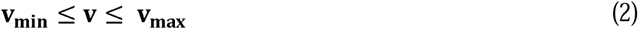

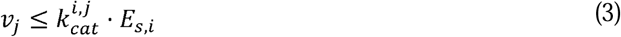

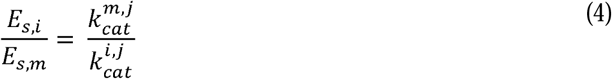

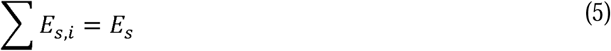

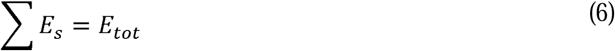

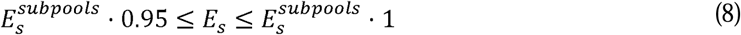

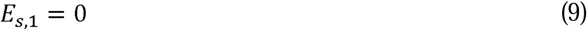

where 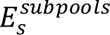 is the enzyme subpool usage for subpool *s* as predicted in the previous step.

Next, we investigated how knocking out the main reaction impacts growth. To this end, we simulated a new round of knockouts starting from the single knockout P4 problem but excluding the constraints on subpools from the previous solutions:

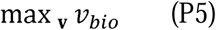

subject to:

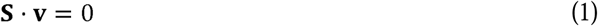

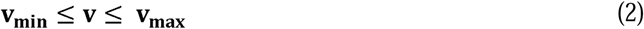

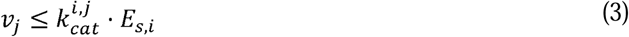

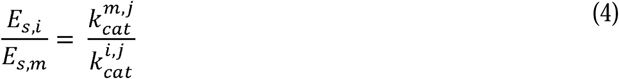

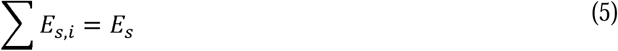

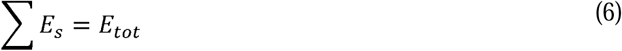

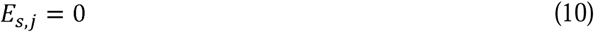

where the subpool *E_s,j_* is the subpool *j* from any enzyme *s*. Then, for all single knockouts that impacted growth, we performed a pairwise double knockout of these subpools as one element of the pair and any other subpool as the other element of the pair to assess how they impact growth:

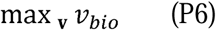

subject to:

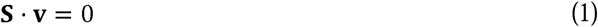

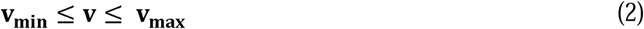

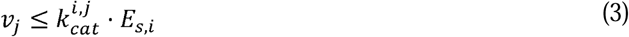

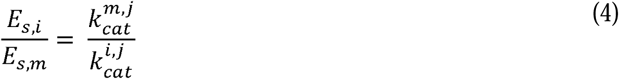

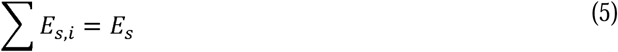

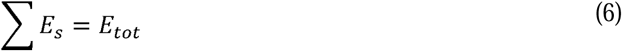

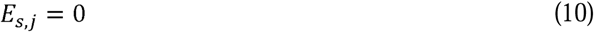

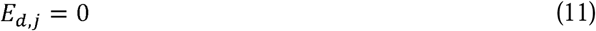

where the subpool *E_d,j_* is the subpool *j* from any enzyme *d* that is not the same as the enzyme *s*. To further assess the effect of knockouts on growth, we performed double knockouts in the conventional GEM, iML1515u, deleting pairs of genes following classical GPR Boolean implementations and optimizing growth as in P6, excluding the enzyme constraints.

Lastly, we were also interested in how knocking out other subpools, besides *E*_*s*,1_, affect growth. We performed a double knockout of subpools from *E*_*s*,2_ to *E*_*s*,5_, since there are few enzymes with more than five subpools.

### 2.5 Flux-sum analysis

To further assess the capabilities of the CORAL model, we investigated the metabolite turnover (flux-sum analysis ^26^) across the optimization problems we solved previously. This ensured that not only we take an enzyme- and reaction-centric overview, but also provide a metabolite-centric assessment. To calculate flux-sums, we apply the following equation:

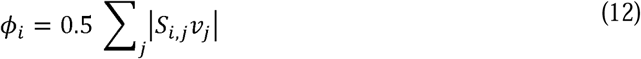

where *ϕ_i_* is the flux-sum of metabolite *i*, *S_i,j_* is the stoichiometric coefficient of metabolite *i* participating in reaction *j*, and *v_j_* is the flux through reaction *j*.

## 3 Results and discussion

### 3.1 CORAL accounts for promiscuous enzyme activity and underground metabolism

By building upon existing protein-constrained approaches, we developed CORAL as a toolbox to investigate promiscuous enzyme activity and underground metabolism in the context of constraint-based modelling. To this end, we first included underground reactions into the *E. coli* iML1515 model, resulting in the iML1515u model. Then, using the DLKcat-predicted *k_cat_* values, we used GECKO 3 to integrate enzyme constraints into iML1515u. The pcGEM was then restructured using CORAL, and the resulting model was named eciML1515u. In CORAL, the enzyme usage in reactions is restructured in a manner that allows for modelling resource usage in promiscuous enzymes. This is achieved by splitting the enzyme pool for each promiscuous enzyme into as many subpools as there are side reactions (Figure 1). The sum of subpools for a certain enzyme then corresponds to the original enzyme pool (see Equation 5, Methods). Before the restructuring, the model contained 3774 metabolites, 8331 reactions, and 1526 enzymes, and application of CORAL resulted in eciML1515u that contains 12048 metabolites, 16605 reactions, 1526 enzymes, and 7260 subpools (Table 1). The large number of metabolites and reactions is due to the added pseudometabolites and pseudoreactions to accommodate the subpools and their usage of the enzyme pools, along with simplification of GPR rules to split enzyme complexes into partial reactions.

**Table 1.** General descriptors for the models used in this study.

|  | **iML1515** | **iML1515u** | **eciML1515u**<br>**(pre-CORAL)** | **eciML1515u**<br>**(CORAL)** |
| --- | --- | --- | --- | --- |
| **Genes** | 1516 | 1530 | 1531 | 1531 |
| **Reactions** | 2712 | 3238 | 8331 | 16605 |
| **Metabolites** | 1877 | 2247 | 3774 | 12048 |
| **Enzymes** | - | - | 1526 | 1526 |
| **Enzyme**<br>**subpools** | - | - | - | 7260 |

### 3.2 Promiscuous enzymes increase metabolic flux variability

Promiscuous enzyme activity and underground metabolism provide alternative flux routes. To investigate the impact of these added reactions, we performed FVA using eciML1515u, both with and without underground reactions. We further tested how chemostat conditions affect flux variability by using a fixed growth rate. We found that flux variability is higher when underground reactions are present (Figure 2A). The scenario without underground reactions and no fixed growth rate had a lower flux variability in 79.85% of reactions when compared to the condition with underground reactions and no fixed growth rate. The scenario with fixed growth rate and without considering underground reactions showed lower flux variability in 79.22% of reactions in comparison to the condition with underground reactions and fixed growth rate. Therefore, we concluded, in line with the expectation, that underground metabolism leads to higher variability of metabolic fluxes.

**Figure 2.**
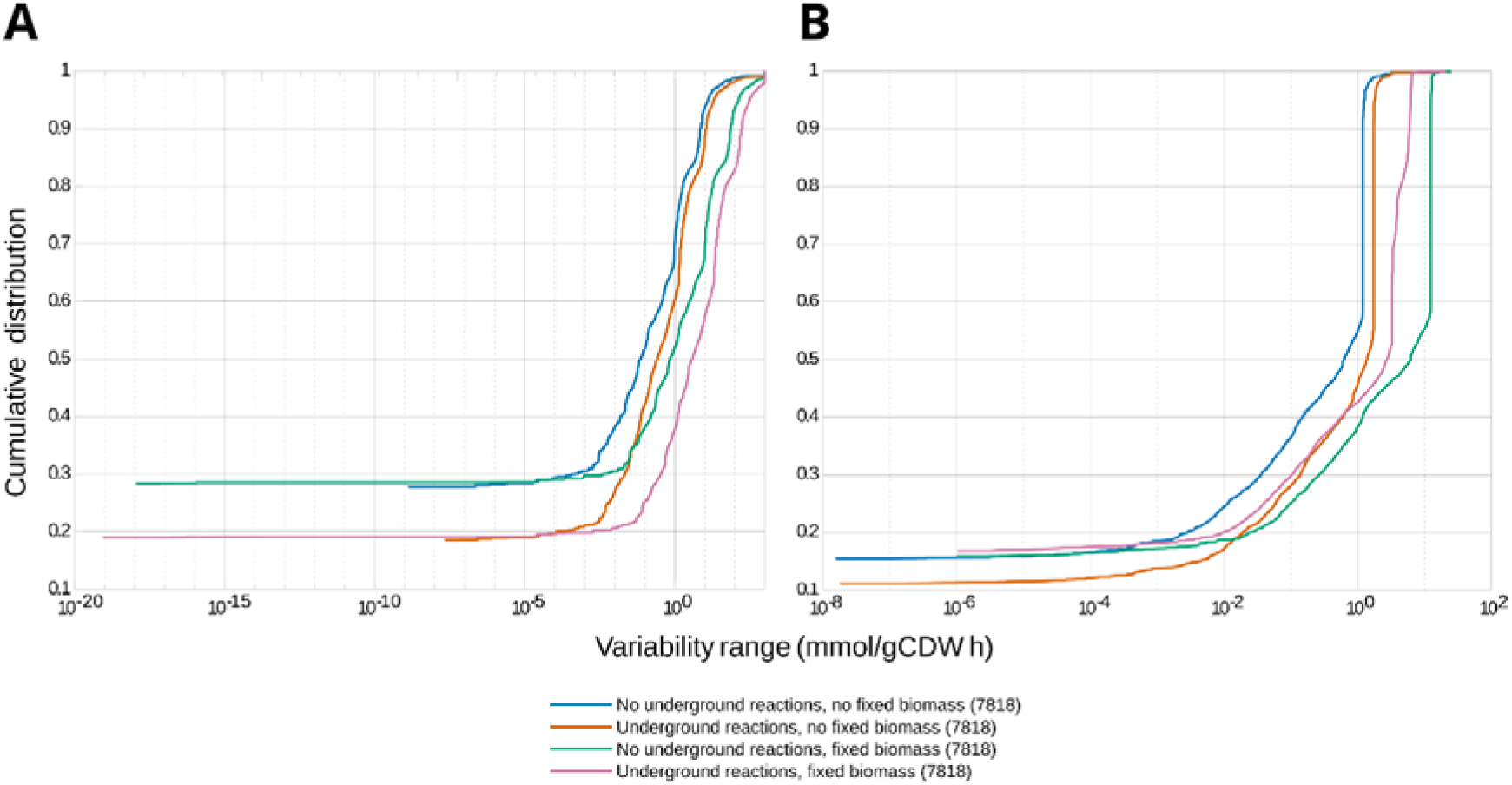
Flux variability for the CORAL-restructured model. We investigated flux and resource allocation variability with or without underground reactions and with or without fixed biomass, as indicated in the legend. **A)** Flux variability for metabolic reactions only (excluding all GECKO- and CORAL-related pseudoreactions). **B)** Flux variability for enzyme subpool usage.

Next, we performed FVA to check the variability of subpool usage instead of metabolic flux. We found that subpool usage variability behaves similarly to metabolic flux variability, with underground reactions leading to an increase in the overall variability (Figure 2B). The subpool usage range was larger in 82.13% of subpools for the condition with underground reactions and no fixed growth rate. Similarly, the subpool usage was larger in 83.30% of subpools for the condition with underground reactions and a fixed growth rate.

### 3.3 Redistribution of enzyme resources to side reactions following knockout of a main reaction ensures metabolic robustness

The restructuring of enzyme usage in CORAL allows for a closer inspection of resource allocation, as promiscuous enzymes are now represented as different pseudometabolites for each reaction it catalyses. Given that, biologically, the larger part of enzyme resources is allocated to the main reaction ^27^, we investigated how these resources are distributed to the side reactions when the main reaction no longer occurs. To this end, we simulated a knockout of the main reaction by setting its enzyme subpool to zero, ensuring that there is no enzyme subpool available to catalyse the main reaction. This is necessary as simulating knockouts using conventional GPR rules would impact all enzyme subpools of a certain enzyme, along with disturbing additional reactions.

In a first round of simulations, we determine the enzyme subpool usage distribution in the wildtype by solving an optimization problem (P3, Methods). Next, we used the enzyme subpool usage distribution to constrain the enzyme subpool usage distributions in the knock-out mutant that abolishes the main activity. These simulations were performed under the same constraints on substrate uptake rates to favour a respiratory metabolism. We found a total of 38 knockout solutions where the main reaction enzyme subpool was used in the wildtype solution and whose knockout resulted in non-lethal mutants.

Considering the changes occurring within an enzyme pool, 13 of the 38 solutions resulted in an almost direct redistribution of *E*_*s*,1_ to *E*_*s*,2_ (which has the second highest catalytic efficiency), such that the enzyme subpool usage value for *E*_*s*,2_ is the same as *E*_*s*,1_ in the P3 solution or within 5% proximity (since we allow for 5% flexibility). For instance, this was the case for the tryptophanase TnaA, whose main subpool catalyses the degradation of tryptophan, but the *E*_*s*,2_ subpool catalyses the degradation of serine. For 10 other solutions, including: TyrB and ArgA, there is a higher allocation to *E*_*s*,2_, while other subpools also receive resources but at a lesser amount. The main subpool of these enzymes catalyses reactions in the biosynthesis of amino acids, as well as the other subpools, although for different amino acids. For seven solutions, including ProC and MtnN, other enzyme subpools received more resources than *E*_*s*,2_. Both are also involved in amino acid biosynthesis, also having side subpools catalysing reactions for different amino acids other than the one in the main reactions. Lastly, in eight solutions, the subpool *E*_*s*,1_ is redistributed entirely to enzyme subpools other than *E*_*s*,2_ (see Supplementary Data 2). For instance, this was the case for TmkK, whose main subpool phosphorylates dTMP to dTDP, and whose other subpools phosphorylates other dNMP nucleotides into dNDPs.

To assess the magnitude of the changes in enzyme subpool allocation after a knockout of the main reaction subpool, we calculated the ratio of how much each enzyme subpool uses from the total enzyme pool (*E_s,j_*/*E_s_*). We found that none of the enzyme subpools that showed a significant increase in the mutant compared to the wildtype were used in the wildtype solution. Inspecting the enzyme subpools with the highest ratio after the knockout, the one with the highest increase belongs to the enzyme pool of the outer membrane porin C, which has 566 subpools. Its main enzyme subpool catalyses the import of calcium in the model. Before the knockout, the main enzyme subpool takes 98.03% of the enzyme pool, whereas after the knockout, the resources were used instead by the subpool 522, which takes up 98.12% of this enzyme pool. The enzyme subpool 522 catalyses the export of L-tryptophan to the environment. Another subpool that significantly takes over the enzyme pool is the subpool 2 of the amino acid acetyltransferase. The subpool 2 catalyses the transfer of the acetyl group from acetyl-CoA into DL-2-aminopimelate, forming 2-acetylaminoheptanedioate and coenzyme A, taking up 83% of the enzyme pool.

Considering the global changes in the model following a knockout, we calculated the difference between enzyme subpool usage predicted for the wildtype and all the mutant solutions. In most solutions, the enzyme subpool usage, with few exceptions, shows a slight change (Figure S1). Among the enzyme subpools with the largest change, there are the enzyme subpools 2 and 3 of the subunit alpha of the ATP synthase complex. Another affected enzyme pool is the subunit beta of the ATP synthase complex, of which the enzyme subpools 2 and 3 also have a higher increase compared to subpools in other enzyme pools. The enzyme subpools 2 and 3 of the ATP synthase gamma chain are also highly impacted. These enzyme subpools are allocated the largest part of the enzyme pool after the knockout of several different main enzyme subpools. These enzyme subpools catalyses the inverse reaction of ATP synthase, producing ADP from ATP hydrolysis. It is important to highlight that these knockouts were not performed *in tandem*, i.e. subunits alpha, beta and gamma are knocked out together, but instead that each subunit of ATP synthase was knocked out individually in separated simulations, which could be a reason why ATP and ADP flux-sums are not impacted (Figure 3). Nonetheless, this result shows that when the main reaction of ATP synthase is blocked, it instead redirects flux to ATP hydrolysis. In *E. coli*, the synthesis or hydrolysis of ATP depends on the direction of mechanical rotation of the ATP synthase complex. This means that CORAL can accurately predict changes in enzyme resource usage for reversible reactions where the reverse reaction has a lower catalytic efficiency.

**Figure 3.**
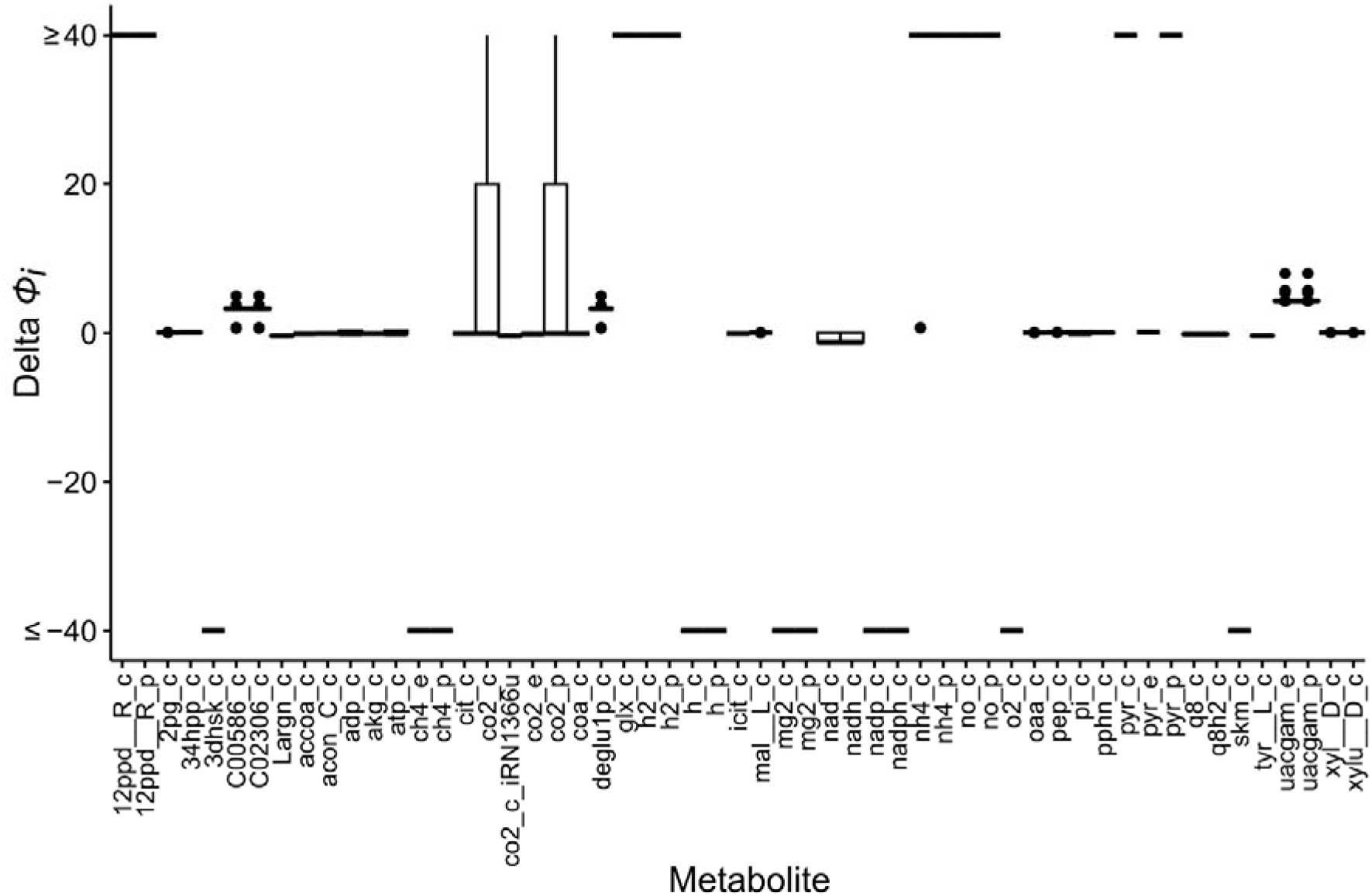
Difference in flux-sums between the wildtype and the mutant solutions. We calculated the flux sums only for metabolites that were common to all solutions when knocking out a single main reaction. We calculated the values by subtracting the flux-sums of the wildtype solution from the flux-sums of the mutant solutions 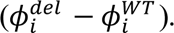 Metabolites with flux-sums of 1000 or -1000 were rescaled to allow better visualization of smaller values. Full names of the metabolites appearing on the x-axis are provided in Supplementary Data 4.

Given that in this analysis the main reaction is knocked out, we inspected if the metabolites involved in these reactions display any change in flux-sums, which can be a useful proxy for metabolite concentrations ^26,28^. We calculated the flux-sums of the wildtype solution and of all 38 mutant solutions and calculated their differences (see Methods). Surprisingly, we found that very few metabolites display changes in flux-sums (Figure 3). Notably, both NAD+/NADH and NADP+/NADPH presented slightly reduced flux-sums, suggesting that in some single knockout solutions, there is a disturbance of redox balance. This can have an impact on central metabolic pathways, since these cofactors are the key electron donors in the respiratory chain and in anabolic reactions. This also leads to oxidative stress and impaired growth and cell survival ^29,30^. In these knockout solutions, however, we do not observe a decrease in growth rate. The metabolite with increased flux-sum is UDP-*N*-acetyl-D-glucosamine, which is the precursor of peptidoglycan and is involved in the biosynthesis of lipopolysaccharides ^31^. Taken together, these results indicate that while a few metabolites are impacted in certain knockout solutions, the cell metabolism is robust to disturbances in enzymes with promiscuous activity, suggesting that having alternative routes can mitigate the impacts of knocking out main reactions.

### 3.4 Double knockouts of pairs of enzymes with no promiscuous activity have notable impact on growth

We next assessed how growth is impacted when pairs of main reactions are blocked. To this end, we knocked out a pairwise combination of all enzyme subpools that catalysed main reactions (e.g., *E*_1,1_ and *E*_2,1_) and compared how it changes compared to a growth rate of 0.11 h^-1^ for the wild type (WT). Altogether, we obtained 3309 pairs out of 24222 whose knockout reduced this growth rate by least 1%. The enzyme subpool pairs used a combination of 156 different enzymes, of which 68 enzymes have promiscuous activity in the model.

Inspecting the knocked-out pairs that most impacted growth (higher than 1% of the growth rate), two enzymes stand out as causing the growth rate to drop no matter the other component of the pair, with the 618 most impacted pairs out of 3309 containing either of these two enzymes. These are the inosine-5’-monophosphate dehydrogenase, whose main reaction in the model catalyses the conversion of inosine-5’-phosphate (IMP) to xanthosine-5’-phosphate (XMP); and the probable acyl-CoA dehydrogenase YdiO, whose main reaction in the model catalyses the conversion of crotonoyl-CoA into butanoyl-CoA using one reduced FAD. Further, knocking out both of these enzymes leads to the largest reduction in the growth rate, with a ratio of growth to that of the wild-type of 0.57. The metabolite xanthosine-5’-phosphate is an intermediary of the *de novo* synthesis of guanine nucleotides, thus being important for growth. Likewise, butanoyl-CoA is an intermediate metabolite in fatty acid metabolism, in both β-oxidation and elongation in mitochondria. In the CORAL model, these enzymes have no promiscuous activity, meaning they have no enzyme subpools to catalyse reactions different than the main reaction.

Amongst the 20912 enzyme pairs that caused no impact on growth (equal or less than 1% of the growth rate), in 15052 pairs there is at least one enzyme with promiscuous activity (Figure 4A), indicating that these have resources redistributed to other enzyme subpools upon knocking out the main reaction enzyme subpool. Meanwhile, in the pairs that affect growth, there is a higher prevalence of enzymes without promiscuous activity (Figure 4B, Supplementary Data 3).

**Figure 4.**
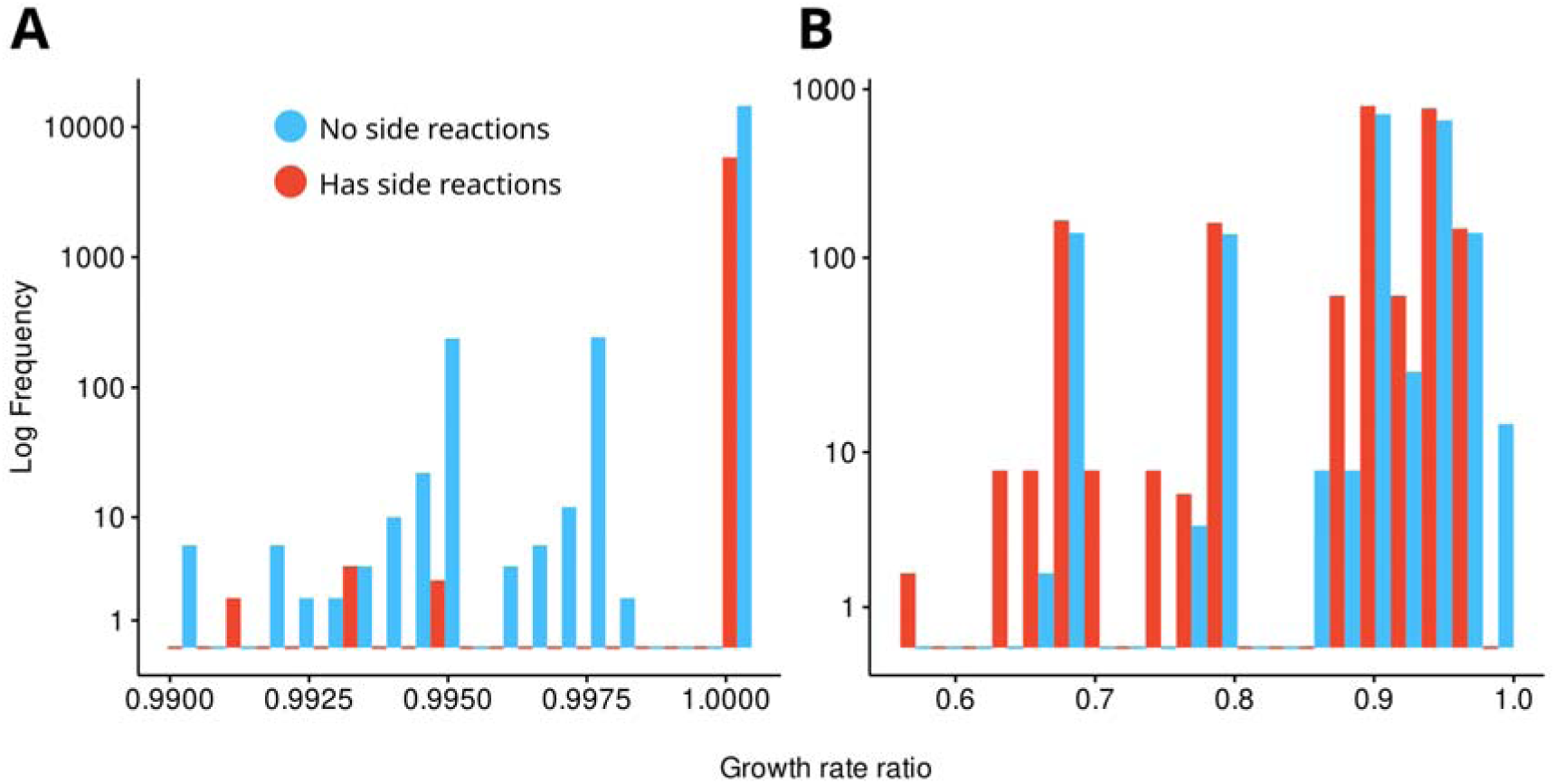
Distribution of growth rate ratios 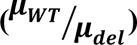 for double knockout mutants. **A)** Growth rate ratios obtained from main reaction pairs that did not affect growth or affected it by less than 1%. **B)** Growth rate ratios obtained from main reaction pairs that affected growth by at least 1%.

To further assess the effects of the double knockouts on growth, we also knocked out pairs of genes from the conventional GEM iML1515u (Figure 5A). We knocked out pairs of genes following GPR rules for all reactions. This is useful to assess how much enzyme promiscuity matters for maintaining the growth rate, since now all reactions controlled by the knocked-out gene will no longer occur. In this scenario, we find that this indeed results in lower growth rates for pairs of genes that code for promiscuous enzymes when no side reactions are available (Figure 5B), suggesting that when all reactions, main or side, catalysed by an enzyme are blocked at the same time, the compensating effect observed in the CORAL model is lost (Figure 5C). Altogether, these results corroborate the findings that promiscuous reactions are important for the maintenance of the growth rate as well as mitigating impacts on cell metabolism.

**Figure 5.**
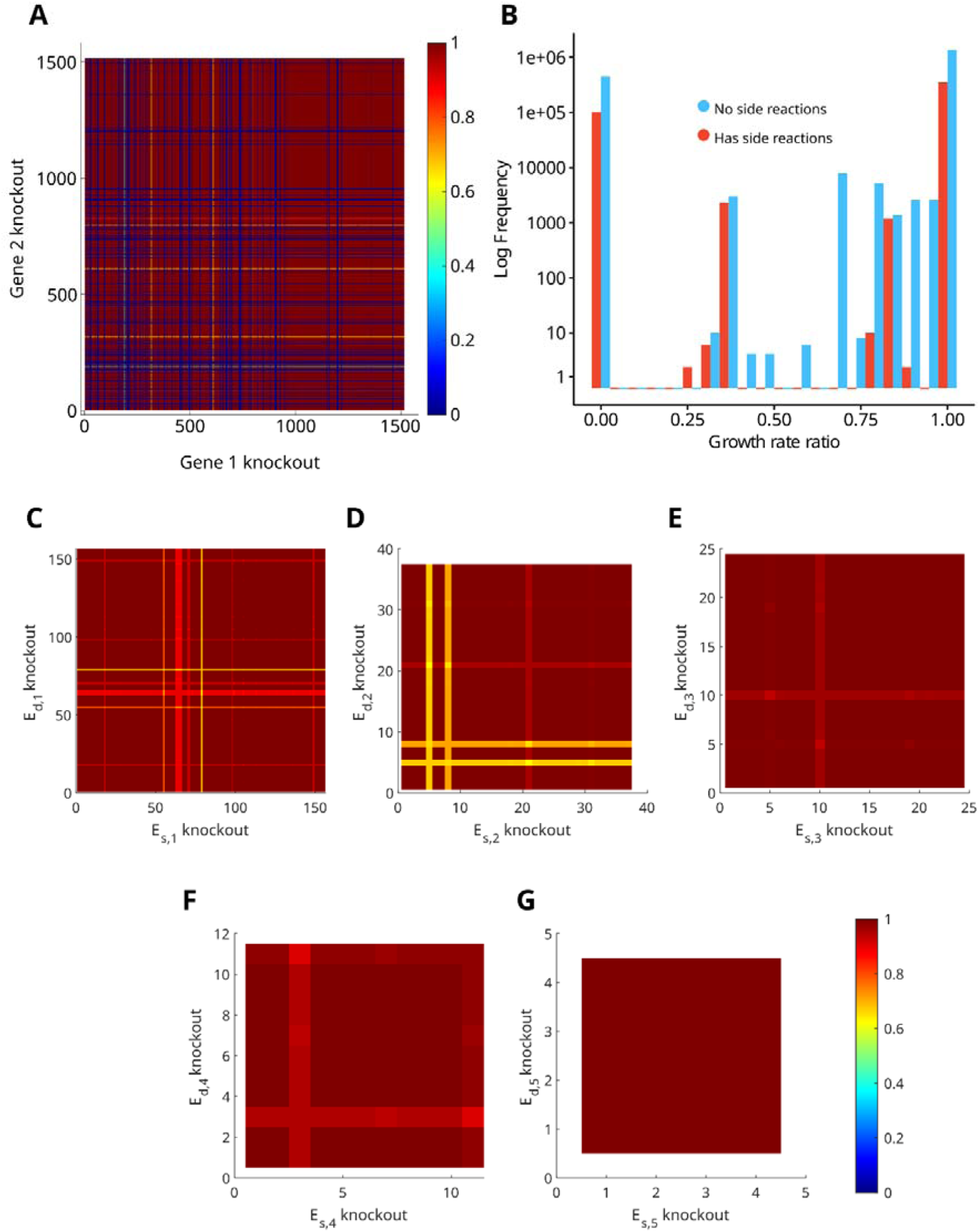
Impact on growth after double knockouts. **A)** Impact on growth following double knockouts in the iML1515u model, performed by deleting pairs of genes according to GPR rules. **B)** Distribution of growth rate ratios following double knockouts in the iML1515u model. **C)** Double knockout of first subpool (*E*_*s*,1_) in the CORAL model. **D)** Second subpool (*E*_*s*,2_). **E)** Third subpool (*E*_*s*,3_). **F)** Fourth subpool (*E*_*s*,4_). **G)** Fifth subpool (*E*_*s*,5_).

We also inspected what happens when we knock out any of the enzyme subpools from *E*_*s*,2_ to *E*_*s*,5_, and noticed that there is little impact on the growth rate. When knocking out the enzyme subpool *E*_*s*,2_ (Figure 5D), there were four enzymes whose presence in a pair had the most impact had the growth rate ratio (≤ 0.71). For the enzyme subpools *E*_*s*,3_ to *E*_*s*,5_, the lowest growth rate ratio was 0.905 (Figure 5E-G). In contrast, knockout pairs of main enzyme subpool had significantly larger impact on the growth rate. This suggests that while side reactions are essential to maintain metabolic and growth balance when the main reaction is impacted, in the opposite situation -- knocking out side reactions while enabling the main reaction -- the double knockouts have negligible impact on growth. This observation holds when growth on different carbon sources are evaluated (Figure S2).

### 3.5 Predictions from CORAL are robust to *k_cat_* values obtained from another predictive tool

The GECKO 3 version has the DLKcat tool integrated into its pcGEM reconstruction pipeline. This has expanded the capabilities of pcGEM reconstruction, given that databases such as BRENDA often have inaccurate or incomplete information, especially for non-model species ^32^. However, DLKcat has been subject to scrutiny given that its predictions of *k_cat_* values are poor for enzymes that differ too much from those in the training dataset ^33^. To assess the robustness of our results, we parameterized the underground reactions of eciML1515u with *k_cat_* values predicted using TurNuP ^23^ and repeated all analyses. We focused on underground reactions since these have non-typical enzyme/substrate pairs and provide a reasonable challenge to *k_cat_* prediction tools.

First, we compare the *k_cat_* values obtained from DLKcat and TurNuP for underground reactions. Interestingly, we found a poor correlation between the two *k_cat_* sets, with a Pearson correlation of 0.21 (Figure S3A). Regarding flux and enzyme usage variability, our results indicated that that the addition of underground reactions still increases the variability in flux and enzyme usage (Figure S3B-C). Regarding the single knockouts of the main reactions, the same metabolic robustness was observed when investigating flux-sums, having most of the same metabolites being impacted (Figure S3D). For the double knockouts that impact growth, we found that the same enzyme subpools are responsible for the decrease in the growth rates as in the DLKcat predicted *k_cat_* values (Figure S3E). Lastly, we also observed that in the knockout pairs that affect growth, there is a higher prevalence of enzymes without promiscuous activity (Figure S3F-G). Taken together, these results show that the predictions obtained with CORAL are robust to the choice of *k_cat_* prediction tool. Given the poor correlation between the two sets of *k_cat_* values, this might be attributed to a compensatory effect by enzyme abundances. Further, considering that native metabolism is governed by reactions catalysed by the main enzyme subpool, changes in side reactions could have limited impact on cell physiology, especially growth.

## 4 Conclusion

Here we present the CORAL Toolbox, a tool for the integration and analysis of promiscuous enzyme activity and underground metabolism. We demonstrated that the inclusion of underground reactions increases metabolic flux variability. Furthermore, the redistribution of enzyme resources from the main to the side reactions highlights the importance of underground metabolism to maintain metabolic flexibility and robustness, serving as alternative routes that compensates for the loss of the main reaction. Moreover, when inspecting the impact of double knockouts of main reactions on growth, promiscuous enzyme activity and underground metabolism proved essential to maintaining growth, as the knockouts with the highest impact were on enzymes without promiscuity. Further, when simulating a scenario where promiscuous enzyme activity does not take place, we observe loss of this compensating effect. Lastly, these results are robust to *k_cat_* values obtained from a different predictive tool.

We note that all of the findings are obtained under the constraint that the ratio of the enzyme allocation to a side reaction in comparison to the corresponding main reaction is reciprocal to the ratio of the respective turnover numbers. This assumption is equivalent to imposing equality between the maximal velocities of the main and side reactions. While, in absence of extensive data on maximal velocities for promiscuous enzymes, this assumption may be plausible, other scenarios can be considered, such as imposing ordering of maximal velocities corresponding to the ordering of turnover numbers. We expect that this relaxation can lead to additional insights about functional implications of promiscuous enzymes.

These results highlight the importance of considering underground metabolism and enzyme promiscuity in terms of metabolic plasticity in constraint-based approaches. The introduced approach allows comprehensive investigations of the interaction between underground metabolism and the rest of the network as well as adaptation to disturbances, pinpointing that flexibility is crucial for cell survival and resilience.

## CRediT authorship contribution statement

**Mauricio Alexander de Moura Ferreira**: Conceptualization, Methodology, Software, Validation, Formal Analysis, Investigation, Writing - Original Draft, Writing - Review & Editing. **Eduardo Luís Menezes de Almeida**: Conceptualization, Methodology, Writing - Review & Editing. **Wendel Batista da Silveira**: Conceptualization, Writing - Review & Editing, Supervision, Project administration, Funding acquisition. **Zoran Nikoloski**: Conceptualization, Methodology, Writing - Original Draft, Writing - Review & Editing, Supervision, Project administration, Funding acquisition.

## Acknowledgements

We thank Dr. Marius Arend, Dr. Philipp Wendering, and Fayaz Soleymani for their support on this study. This study was financed in part by the Coordenação de Aperfeiçoamento de Pessoal de Nível Superior – Brasil (CAPES) – Finance Code 001.

## Data availability

The CORAL Toolbox is publicly available in a GitHub repository along with the code and data for reproducing this work: https://github.com/mauricioamf/CORAL.

## Supplementary figures

**Figure S1.**
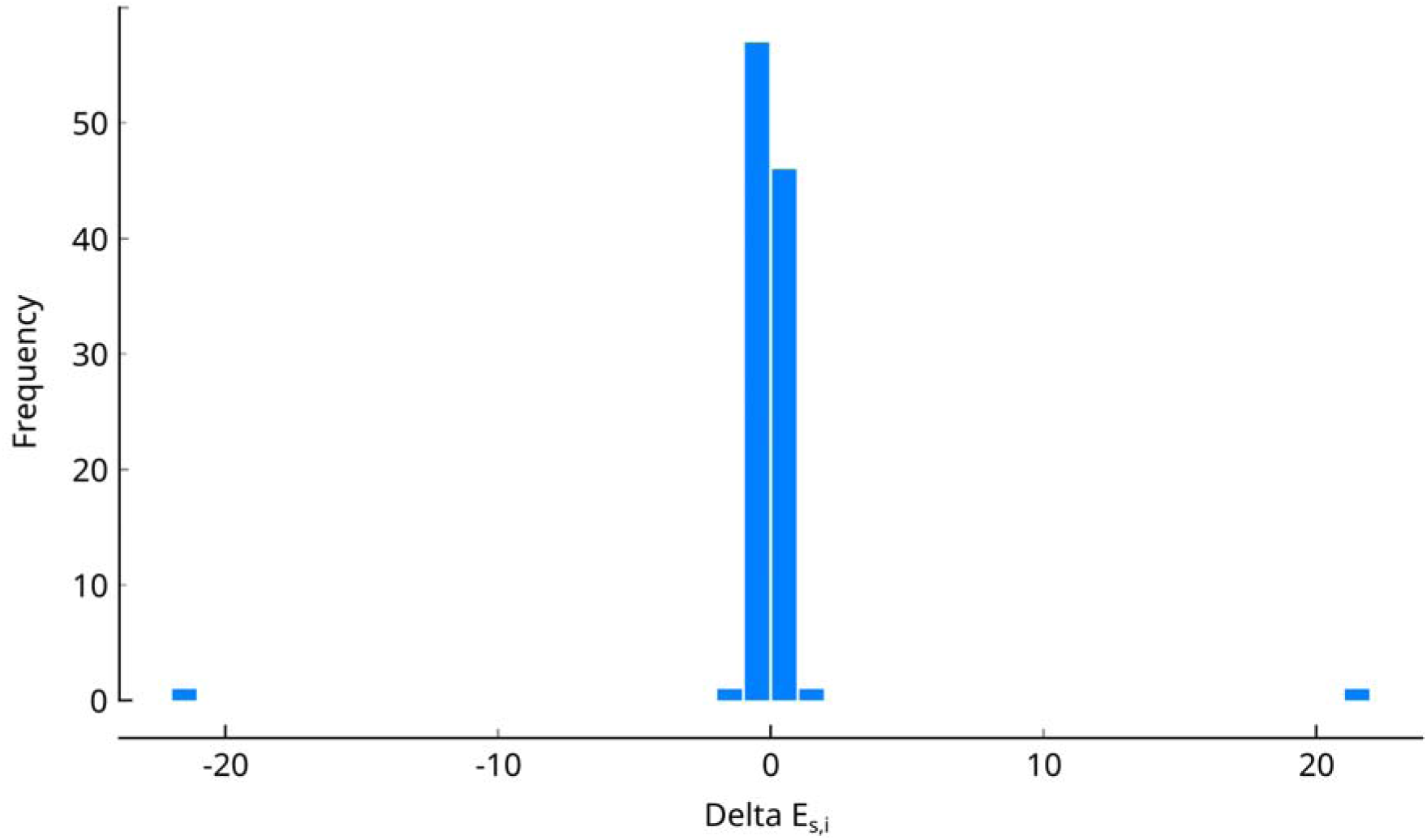
Changes in enzyme subpool usage following the knockout of a single main reaction. We calculated delta by subtracting the enzyme subpool usage of the wildtype from the enzyme subpool usage of the mutant solutions.

**Figure S2.**
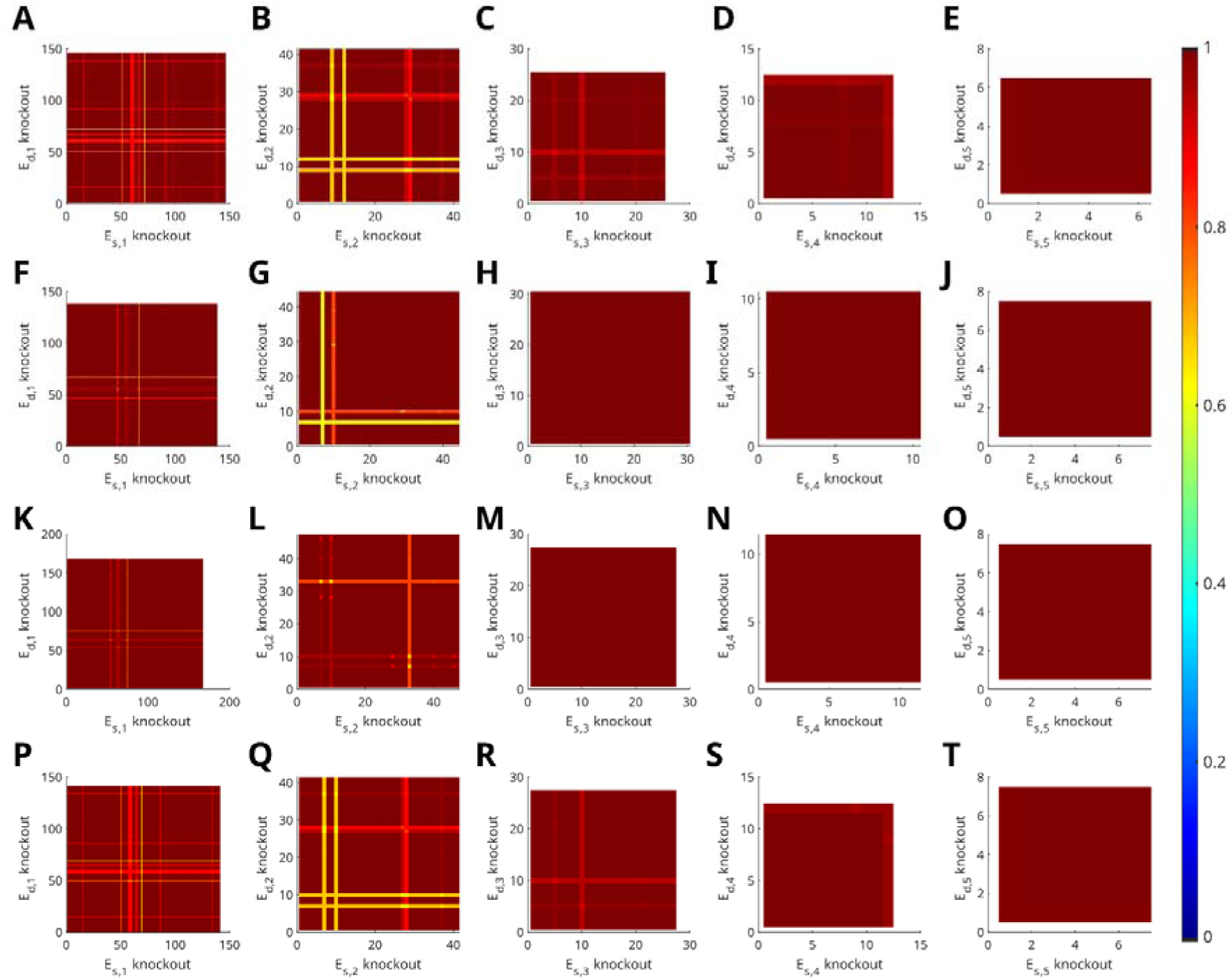
Impact on growth after double knockouts considering alternative carbon sources. **A)** Impact on growth after a double knockout of first subpool (*E*_*s*,1_) using arabinose as carbon source. **B)** Second subpool (*E*_*s*,2_). **C)** Third subpool (*E*_*s*,3_). **D)** Fourth subpool (*E*_*s*,4_). **E)** Fifth subpool (*E*_*s*,5_). **F)** Impact on growth after a double knockout of first subpool (*E*_*s*,1_) using fructose as carbon source. **G)** Second subpool (*E*_*s*,2_). **H)** Third subpool (*E*_*s*,3_). **I)** Fourth subpool (*E*_*s*,4_). **J)** Fifth subpool (*E*_*s*,5_). **K)** Impact on growth after a double knockout of first subpool (*E*_*s*,1_) using fucose as carbon source. **L)** Second subpool (*E*_*s*,2_). **M)** Third subpool (*E*_*s*,3_). **N)** Fourth subpool (*E*_*s*,4_). **O)** Fifth subpool (*E*_*s*,5_). **P)** Impact on growth after a double knockout of first subpool (*E*_*s*,1_) using xylose as carbon source. **Q)** Second subpool (*E*_*s*,2_). **R)** Third subpool (*E*_*s*,3_). **S)** Fourth subpool (*E*_*s*,4_). **T)** Fifth subpool (*E*_*s*,5_).

**Figure S3.**
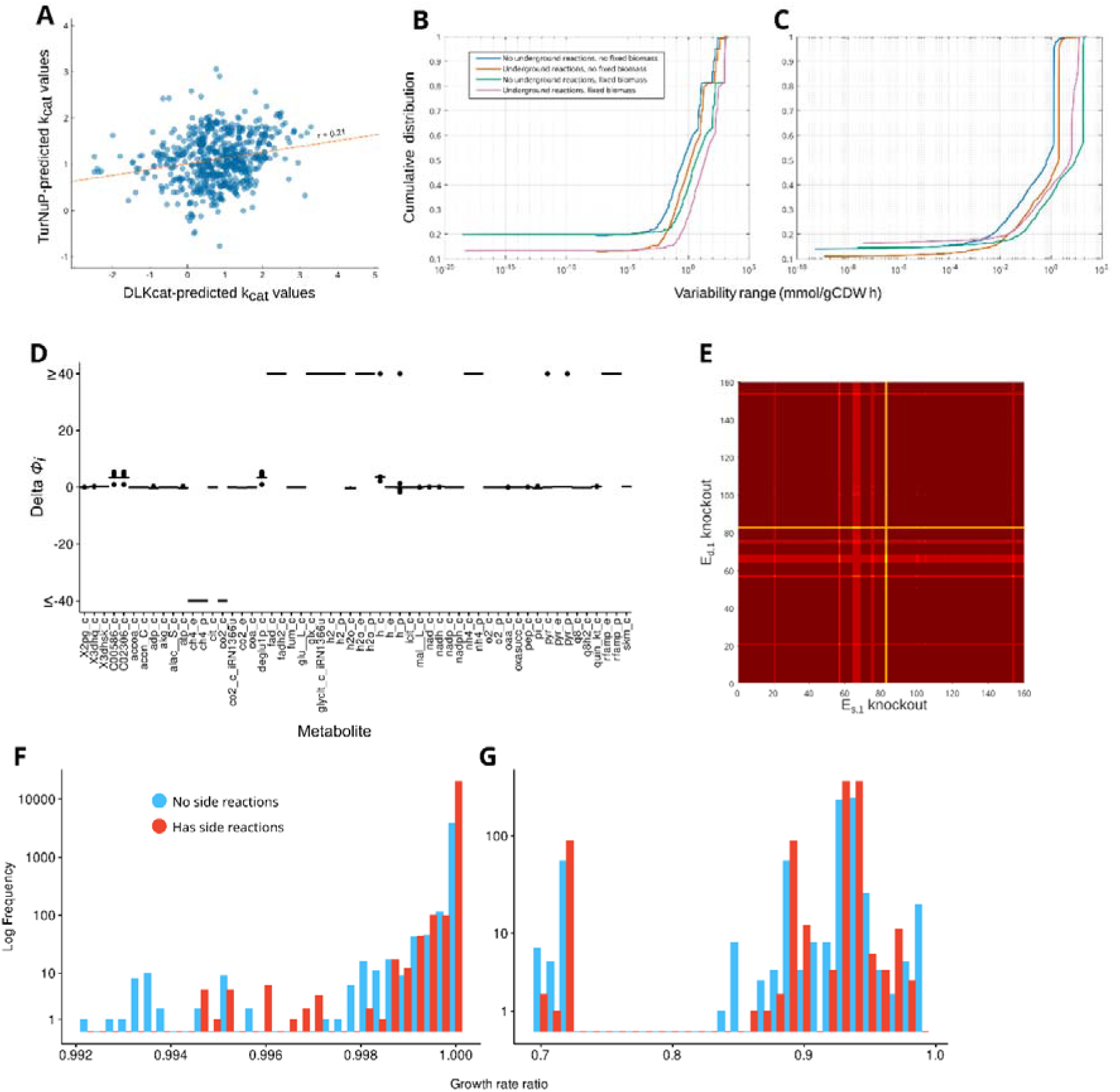
Robustness analysis of CORAL using TurNuP-predicted *k_cat_* values. We performed all analyses as done when using DLKcat-predicted *k_cat_* values. **A)** Comparison of DLKcat- and TurNuP-predicted *k_cat_* values. **B)** Flux variability for metabolic reactions only (excluding all GECKO- and CORAL-related pseudoreactions). **C)** Flux variability for enzyme subpool usage. D) Difference in flux-sums between the wildtype and the mutant solutions. Full names of the metabolites appearing on the x-axis are provided in Supplementary Data 4. **E)** Impact on growth after double knockouts. **F)** Growth rate ratios obtained from main reaction pairs that did not affect growth or affected it by less than 1%. **G)** Growth rate ratios obtained from main reaction pairs that affected growth by at least 1%.

## Description of Additional Supplementary Files

**File Name: Supplementary Data 1**

Description: List of added reactions and their corresponding *k_cat_* values, predicted from either DLKcat or TurNiP.

**File Name: Supplementary Data 2**

Description: List of subpools and their corresponding percentage of the enzyme pool.

**File Name: Supplementary Data 3**

Description: List of main subpool knockout pairs and their corresponding growth rate before and after knockout.

**File Name: Supplementary Data 4**

Description: List of metabolites evaluated during flux-sum analysis of single knockout solutions for both DLKcat- and TurNuP- parameterized models.

