## Supplementary text for "Integrating promiscuous enzyme activities in protein-constrained models pinpoints the role of underground metabolism in robustness of metabolic phenotypes"

### Supplementary methods

The CORAL Toolbox introduces a simplification of GPR rules by splitting all reactions catalysed by enzyme complexes (“AND” rules) into multiple partial reactions catalysed by one enzyme each. Consider the example reaction:

substrate1[c] + substrate2[c] + enzyme1[c] + enzyme2[c] + enzyme3[c] => product1[c]

As it needs three different enzymes, it is split into three different reactions. In the first reaction, the product of the original reaction is replaced by a pseudometabolite, and contains only the first enzyme, enzyme1. The second reaction uses the enzyme enzyme2 and takes pseudometabolite1as the substrate, and produces the second pseudometabolite, pseudometabolite2. The third reaction uses the enzyme enzyme3, consumes pseudometabolite2, and yields the product of the original reaction, product1:

substrate1[c] + substrate2[c] + enzyme1[c] => pseudometabolite1[c]

pseudometabolite1[c] + enzyme2[c] => pseudometabolite2[c]

pseudometabolite2[c] + enzyme3[c] => product1[c]

The GECKO 3 formulation of pcGEMs directly represents enzymes in the stoichiometric matrix, where each enzyme corresponds to a pseudometabolite that participates in a reaction, with a pseudo-stoichiometric coefficient given by the ratio between the molecular weight of the enzyme (MW) and its turnover number, $k_{cat}$, for the reaction. In this formulation, different reactions can be catalysed by the same enzyme. This can be exemplified as:

substrate1[c] + substrate2[c] + enzyme1[c] => product1[c] + product2[c]

substrate3[c] + substrate4[c] + enzyme1[c] => product3[c] + product4[c]

enzyme1[c] => prot_pool[c]

prot_pool[c] =>

In the CORAL reformulation, promiscuous enzymes are accounted for by splitting the pool of an enzyme that catalyses more than one reaction into multiple subpools, with each subpool responsible for the enzyme resources of only one reaction. In the model, the example reactions from above are reformulated as:

substrate1[c] + substratS2[c] + enzyme1_1[c] => product1[c] + product2[c]

substrate3[c] + substrate4[c] + enzyme1_2[c] => product3[c] + product4[c]

enzyme1_1[c] => pool_enzyme1[c]

enzyme1_2[c] => pool_enzyme1[c]

pool_enzyme1[c] => prot_pool[c]

prot_pool[c] =>

where enzyme1_1 and enzyme1_2 are treated as different pseudometabolites.
